# Human fetal neural stem cell-derived astrocytes maintain glutamate transport after hypoxic injury *in vitro*

**DOI:** 10.1101/2021.05.17.444180

**Authors:** Vadanya Shrivastava, Devanjan Dey, Chitra Mohinder Singh Singal, Paritosh Jaiswal, Ankit Singh, JB Sharma, Parthaprasad Chattopadhyay, Jayanth Kumar Palanichamy, Subrata Sinha, Pankaj Seth, Sudip Sen

## Abstract

Astrocytes are the most abundant glial cells that play many critical roles in the central nervous system physiology including the uptake of excess glutamate from the synapse by Excitatory Amino Acid Transporters (EAATs). Among the EAATs, EAAT2 are predominantly functional, astrocyte-specific glutamate transporters in the forebrain. Hypoxic brain injury is a pathological phenomenon seen in various clinical conditions including stroke and neonatal hypoxic ischemic encephalopathy. Glutamate excitotoxicity is an important cause of neuronal cell death in disorders involving hypoxic brain injury. As findings from rodent models cannot always be reliably extrapolated to humans, we aimed to develop a homogenous population of primary human astrocytes to study the effect of hypoxic injury on astrocyte function, especially glutamate uptake. We successfully isolated, established and characterized cultures of human fetal neural stem cells (FNSCs) from aborted fetal brains. FNSCs were differentiated into astrocytes, and characterized by increased expression of the astrocyte markers, glial fibrillary acidic protein (GFAP), EAAT1 and EAAT2. A concomitant decrease in neural stem cell marker, Nestin, was observed. Differentiated astrocytes were exposed to various oxygen concentrations mimicking normoxia (20% and 6%), moderate and severe hypoxia (2% and 0.2% respectively). Interestingly, no change was observed in the expression of glutamate transporter, EAAT2 and glutamate uptake by astrocytes, even after exposure to hypoxia. Our novel model of human FNSC derived astrocytes exposed to hypoxic injury, establishes that astrocytes are able to maintain glutamate uptake even after exposure to severe hypoxia for 48 hours, and thus provides evidence for the neuroprotective role of astrocytes in hypoxic injury.

## Introduction

The central nervous system comprises of different cell types including neurons and non-neuronal glial cells. Astrocytes, conventionally considered as supporting glial cells of the brain, also account for other important roles like providing energy intermediates to neurons, maintaining the water and ionic balance in and around them, formation and regulation of the blood-brain barrier, calcium signaling, along with release and uptake of neurotransmitters, especially that of glutamate and GABA (Chandrasekharan et al., 2016). Glutamate is an important excitatory neurotransmitter in the brain and astrocytes play a key role in preventing glutamate excitotoxicity by taking up excess glutamate from the synapse via the excitatory amino acid transporters (EAATs).

Hypoxic injury to brain cells leads to neuronal cell death and results in debilitating motor and cognitive disabilities. Excitotoxicity is one of the major mechanisms of neuronal damage in hypoxic brain injury (Fujikawa, 2015). Given the crucial role of astrocytes in maintaining the extracellular glutamate levels, their dysfunction in hypoxia can adversely affect neuronal survival following hypoxic injury. Earlier studies investigating astrocyte glutamate dynamics in hypoxia have used rodent model systems to prove their point. While these models give important insights into the mammalian brain, fundamental differences exist between the rodent and human brain, which differ not only in size and number of neurons but also in morphological and functional diversity, as well as gene expression and its regulation (Hodge et al., 2019). Therefore, there is a need to study the behavior of human neural cell types in hypoxic injury.

This study aimed to evaluate the role of astrocytes in hypoxic injury by developing an *in vitro* model, comprising of human astrocytes derived from fetal neural stem cells. Using this model system, the effect of hypoxia on astrocyte function was evaluated, particularly the expression and function of glutamate transporter.

## Methodology

### Sample collection

Aborted fetal samples were collected (n=5), after taking informed consent from mothers undergoing Medical Termination of Pregnancy (MTP) in the Department of Obstetrics and Gynecology, AIIMS, New Delhi, India. Necessary approvals were taken from Institutional Ethics Committee and Institutional Committee for Stem Cell Research, before starting the study and the study was carried out in conformation with the World Medical Association Declaration of Helsinki. Fetal samples from mothers undergoing MTP in the second trimester of pregnancy (12-20 weeks) for maternal indications, were included, while fetal indications (such as chromosomal anomalies) were excluded from the study.

### Isolation of human fetal neural stem cells (FNSCs)

Isolation of human FNSCs from aborted fetuses was done in accordance with previous protocols (Tewari et al., 2015). Briefly, hNSCs were isolated from the subventricular zone of the fetal brain and subsequently plated onto poly-D-lysine coated culture flasks in neural stem cell media containing Neurobasal media (GIBCO, NY, USA) with 1% N2 supplement (GIBCO, NY, USA), 2% Neural survival factor-1 (Lonza, IA, USA), 1% Glutamax (GIBCO, NY, USA), 5mg/mL of bovine serum albumin (Sigma, MO, USA), penicillin (50 IU/ml), streptomycin (50 µg/ml) and gentamicin (2 µg/ml). Neuro-spheres were observed after 2-3 days, dissociated by gentle agitation during sub-culture, and then seeded onto poly-D-lysine coated flasks to allow their adherence, to generate monolayers of FNSCs. Differentiation into astrocytes was initiated after 2-3 passages.

### Differentiation of human fetal neural stem cells into astrocytes

FNSC growth media was substituted by Minimal essential media (MEM) (Sigma, MO, USA) with 10% FBS to induce differentiation of FNSCs into astrocytes (Tewari et al., 2015). Half the media was changed every alternate day for 14 days, and sub-culturing was done when cells reached 80% confluency. Astrocytes were characterized by presence of GFAP, EAAT1 and EAAT2 in cells.

### Exposure of differentiated human astrocytes to different oxygen concentrations

Differentiated human astrocytes (at day14), at 70-80% confluency, were exposed to oxygen concentrations mimicking normoxia (20% and 6%) and hypoxia (2% and 0.2%) for 48 hours, at 37°C and 5% CO_2_ that was created using an Anoxomat hypoxia induction system (Advanced Instruments, Norwood, MA, USA). Hypoxia exposure was validated by evaluating the expression of HIF1α and hypoxia-responsive genes CA9, VEGF and PGK-1.

### RNA isolation, cDNA synthesis and qPCR

Total RNA was extracted from the cells at different stages of differentiation after exposure to various oxygen concentrations, using Tri-Reagent (Sigma, MO, USA) and quantified by Nano-Drop ND-1000 spectrophotometer (Thermo-Fisher Scientific, MA, USA). cDNA was synthesized with 1µg total RNA using M-MuLV-RT (Thermo-Fisher Scientific, MA, USA) and random hexamer primers (IDT, IL, USA) as described previously (Dey et al., 2020). The expression of various genes was evaluated in the cells (in triplicates) using gene-specific primers (IDT, IL, USA) (Table 1) and DyNAmo Flash SYBR Green qPCR kit (Thermo-Fisher Scientific, MA, USA) using CFX96 Touch™ Real-Time PCR Detection System (BioRad, CA, USA). 18S rRNA was used as an internal reference gene for normalization. Relative fold change in gene expression was calculated using 2^-ΔΔCT^ method. For differentiation experiments, day 0 samples (human FNSCs) were used as controls, while differentiated astrocytes (day 14) exposed to 20% oxygen (mimicking normoxia) were the controls for hypoxia experiments.

**Table 1:**
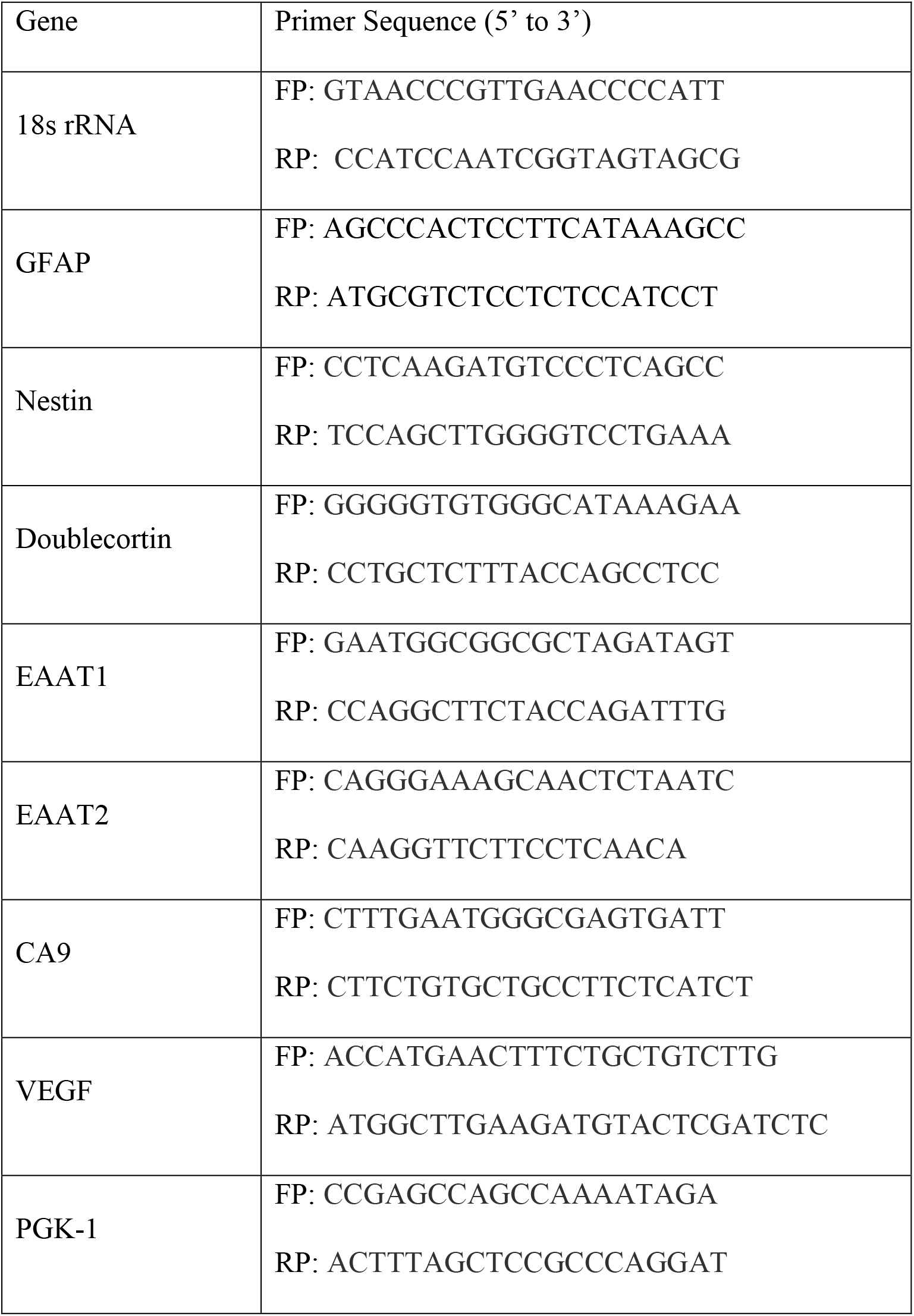
Sequences of primers used for gene expression analysis (FP = Forward primer; RP = Reverse primer)

### Western blotting

Protein was isolated using Tri-Reagent (Sigma, MO, USA) as per the manufacturer’s protocol and quantified by BCA method (Thermo-Fisher Scientific, MA, USA). Equal amounts of protein extracts (20µg) were electrophoresed on 10 – 15% SDS-polyacrylamide gels and electro-transferred onto nitrocellulose membrane (BioRad, Hercules, CA, USA). The membrane was blocked with 5% non-fat milk (NFM) dissolved in 0.1% Tween-20 containing tris-buffered saline containing (TTBS) for 1 hour. The blots were probed overnight with specific primary antibodies (Table 2), that were diluted in 1% NFM in TTBS. After washing with TTBS, the membrane was incubated for 1 hour with the appropriate secondary antibody (Table 2). The blot was incubated with Luminol and peroxidase (Abbkine SuperLumia ECL Plus Kit, Hubei, China) and chemiluminescence detection was done using Azure Biosystems c280 gel documentation sytem (Dublin, CA, USA), followed by analysis with Image J software. Normalization was done using GAPDH and β-actin.

**Table 2:** Specifications for primary and secondary antibodies used for western blot experiments.

| Antibody | Molecular weight | Antibody specifications | Antibody dilution | Procured from |
| --- | --- | --- | --- | --- |
| Anti-GFAP | 50 kDa | Polyclonal<br>Rabbit IgG | 1:1000 | Abbkine,<br>China |
| Anti-EAAT2 | 60 kDa | Polyclonal<br>Rabbit IgG | 1:1000 | Abbkine,<br>China |
| Anti- HIF1α | 120 kDa | Monoclonal<br>Rabbit IgG | 1:1000 | CST, MA,<br>USA |
| Anti-GAPDH<br>(loading control<br>for differentiation<br>experiments) | 37 kDa | Monoclonal<br>Mouse IgG | 1:2000 | Abgenex,<br>India |
| Anti β-actin<br>(loading control<br>for hypoxia<br>experiments) | 42 kDa | Polyclonal<br>Rabbit IgG | 1:1000 | St. John's<br>Laboratory,<br>UK |
| HRP-tagged Anti-rabbit secondary<br>antibody | - | Goat Anti-rabbit IgG | 1:2000 | CST, MA,<br>USA |
| HRP-tagged anti-<br>mouse secondary<br>antibody | - | Horse Anti-<br>mouse IgG | 1:2000 | CST, MA,<br>USA |

### Flow cytometry

Cells (at different stages of differentiation) were fixed using 2% paraformaldehyde and then permeabilized with 1% BSA containing 0.1% Triton X-100. Cells were then blocked in 2% BSA for half an hour and subsequently stained (intracytoplasmic) with Alexa Fluor 647 conjugated rabbit anti-human GFAP antibody (BD Biosciences, cat. no. 561470) using appropriate controls. Cells were washed, resuspended in 2% paraformaldehyde, and data was acquired using BD LSR Fortessa (BD Biosciences, San Jose, CA, USA) and analyzed using FlowJo v10 software.

### Immunocytochemistry

Cells were plated onto coverslips, washed once with PBS and fixed with 4% PFA. The cells were then incubated for 1 hour in 1% BSA with 0.5% Triton X-100 and then washed with PBS before incubating overnight at 4°C with primary antibody (Rabbit Anti-Nestin 1:1000 (Millipore, Cat no. ABD69). Thereafter, cells were washed thrice with PBS and then incubated for an hour in secondary antibody (Mouse anti-Rabbit FITC 1:1000 (Invitrogen, Cat no. A11008) for 1 hour at room temperature. Cells were then washed thrice with PBS, and mounted onto glass slides using Vectashield mountant containing DAPI. The slide was allowed to dry overnight. Images were taken on Nikon Eclipse Ti-S fluorescent microscope (Tokyo, Japan) and analyzed with NIS-Elements BR 4.30.00 64-bit software.

### Glutamate uptake assay

Glutamate uptake by astrocytes was measured as described by Lutgen et al. (2016) with minor modifications. Briefly, after exposing astrocytes to hypoxia for 48 hours, supernatant in culture flasks were discarded and replaced with 2 mL of HBSS (ThermoFisher) with 2mM monosodium glutamate. Cells were then incubated for 30 min at 37°C in normoxic conditions (20% O_2_), following which, the supernatant was removed and snap-frozen for estimation of glutamate, while the cells were treated with TRIzol for isolation of RNA and protein. Glutamate was estimated in the supernatant by a fluorimetric assay (Abcam cat.no. ab138883) which used a glutamate dehydrogenase coupled mechanism for estimation. Glutamate concentrations in 2 mL HBSS containing 2 mM glutamate (vehicle control) were estimated and found to be ∼2.06 mM. Glutamate uptake values were normalized to the amount of protein obtained in the corresponding cells (n=7).

### Statistical analysis

Statistical analysis was done using Graph Pad Prism v6. Kruskal Wallis test was used to detect significant differences between groups. In datasets that showed a significant p-value on the Kruskal-Wallis test, a comparison between specific groups was made by Dunn’s pairwise comparison test. p-value < 0.05 was considered statistically significant.

## Results

### Isolation and characterization of human fetal neural stem cells

Neural stem cells were isolated from the subventricular zone of the fetal brain (n=5). Neurospheres were observed after 48-72 hrs in culture which showed FNSCs radiating outward from the core (Fig. 1A). Neurospheres were dissociated and sub-cultured, to form monolayers. FNSCs cultured in monolayer appeared as small cells with unipolar morphology (Fig. 1B). Human FNSCs were expanded and characterized by the expression of Nestin on immunocytochemical staining (Fig. 1C).

**Figure 1.**
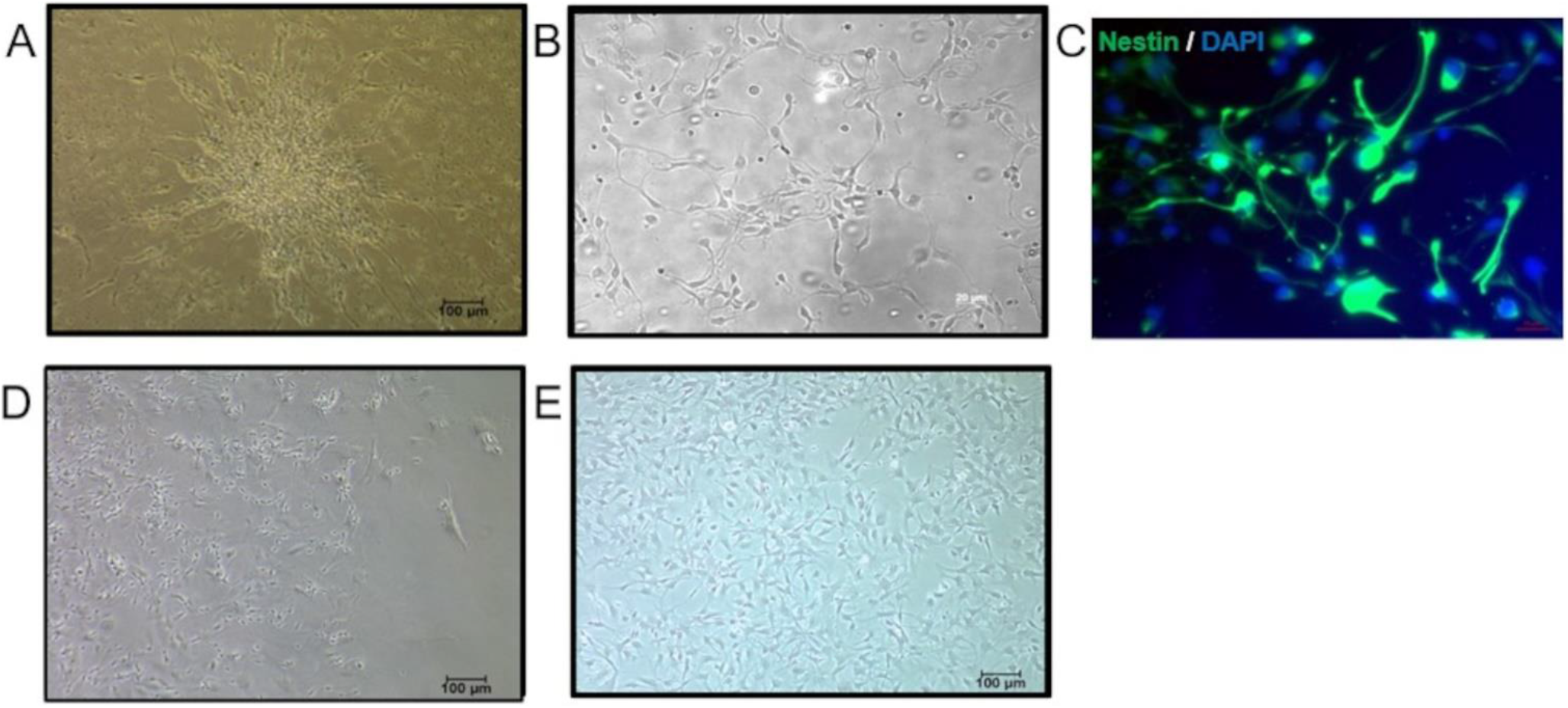
Morphological features of human fetal neural stem cells and differentiating astrocytes. (A) Neurosphere formation seen after 24-48 hours of FNSC isolation. (B) FNSCs in monolayer culture display unipolar morphology. (C) Immunofluorescence image of FNSCs stained for neural stem cell marker Nestin (green). Nuclei are stained with DAPI (blue). Morphological changes seen at (D) day 7, (E) day 14 of astrocytic differentiation. Differentiated cells are larger and show flattened morphology.

### Differentiation of human FNSCs into astrocytes

After 3-4 passages, FNSCs were differentiated into astrocytes (n=5) by replacing the neural stem cell media with complete MEM media, and differentiation was monitored over 14 days. With the progress of differentiation, cells were observed to change morphology from small unipolar cells to large, flat cells with large nuclei (Fig. 1D and 1E). Cells were harvested at days 0, 7 and 14 to evaluate change in expression of specific markers.

### Expression of astrocytic markers in differentiating cells

There was an increase in the mRNA expression of the astrocyte-specific marker GFAP from day 0 to day 7 (fold change 4.59 ± 2.48, p-value > 0.05) and day 14 (fold change 8.6±5, *p* = 0.0079) (Fig. 2A). These changes at the mRNA level were mirrored in GFAP protein expression by western blotting (Fig. 2C). An increase was observed in the normalized expression of GFAP from 0.28 ± 0.17 at day 0, to 1.03 ± 0.2 at day 7 (*p* > 0.05), and 1.55 ± 0.05 at day 14 (*p* = 0.0134), equating to a fold change of 7.2 ± 4.4 at day 14 (Fig. 2D). Differentiating cells also showed an increase in the mRNA expression of astrocyte specific glutamate transporters (Fig. 2B). EAAT1 expression increased with a fold change of 1.4 ± 0.3 (p >0.05) at day 7 and fold change of 2.90 ± 0.67 at day 14 (p = 0.02). EAAT2 also increased over the course of differentiation, with fold changes of 1.1 ± 0.08 (p > 0.05) at day 7 and fold change of 2.93 ± 1.9 (p = 0.019) at day 14 of differentiation. Differentiating cells (day 0 and day 14) were also analyzed for the expression of GFAP, by flow cytometry (Fig. 3A – D). The percentage of cells expressing GFAP increased significantly from 36 ± 7.8 percent (day 0) to 94.97 ± 1.7 percent (day 14) (*p*=0.03) (Fig. 3E).

**Figure 2.**
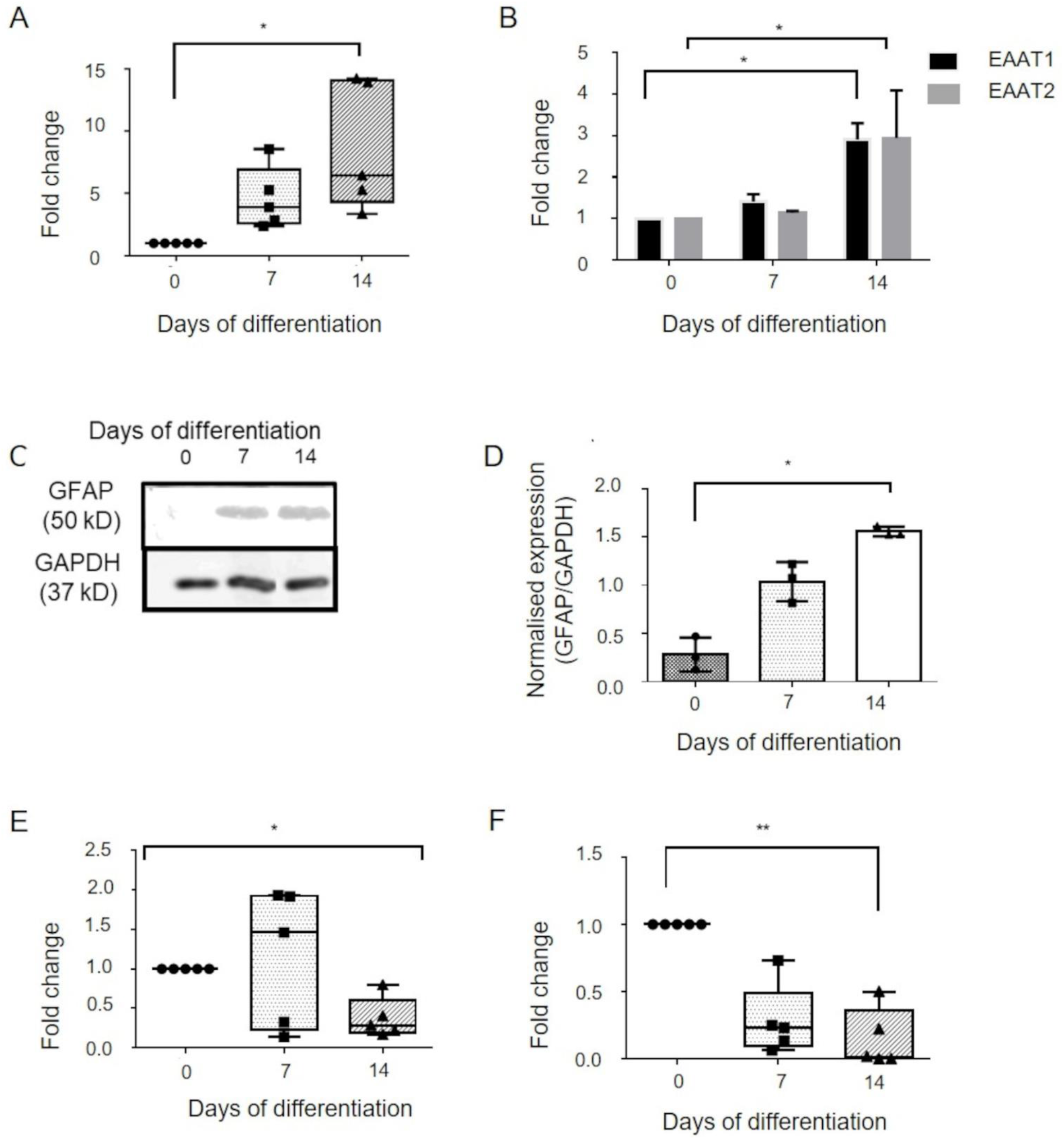
Expression of lineage markers during differentiation of human FNSCs into astrocytes. Expression of astrocytic marker (A) GFAP (n=5), (B) EAAT1 and EAAT2 (n=3) at various days of differentiation as obtained on qPCR analysis. (C) Representative western blot image showing protein expression of GFAP at various days of differentiation. GAPDH is loading control. (D) Quantification of western blot data (n=3). Expression of neural stem cell marker Nestin (E) and neuronal marker Doublecortin (F) at various days of differentiation as obtained on qPCR analysis (n=5). qPCR data is represented as fold change in gene expression obtained using day 0 samples as control. GFAP: Glial fibrillary acidic protein. EAAT1: Excitatory amino acid transporter 1. EAAT2: Excitatory amino acid transporter 2. GAPDH: Glyceraldehyde 3-phosphate dehydrogenase. Data represented as Mean±SD. * p-value <0.05 **p-value <0.01

**Figure 3.**
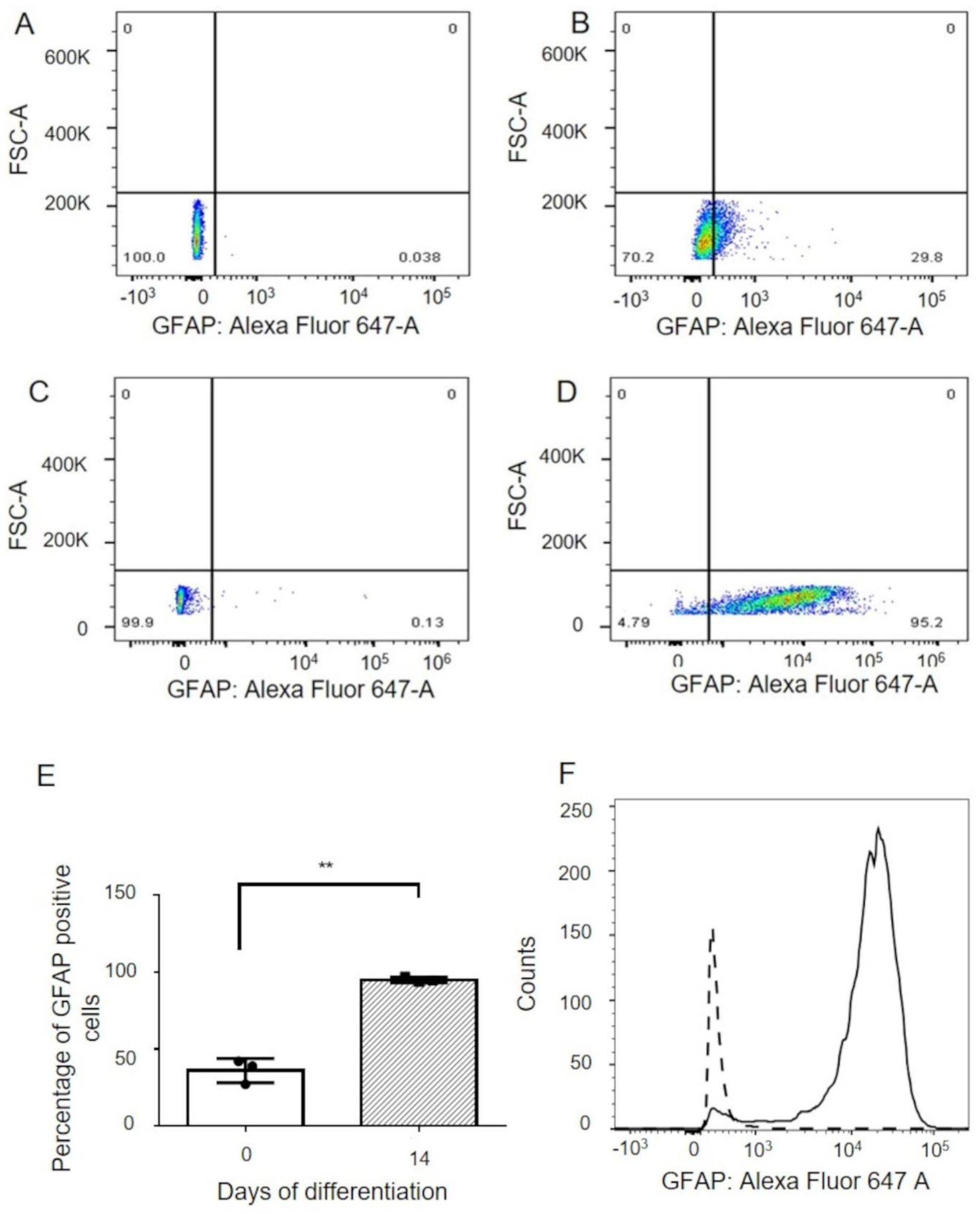
Flow cytometric analysis of GFAP expression during astrocytic differentiation. Dot plots showing (A) Unstained and (B) stained populations of human FNSCs (day 0). Representative dot plots for (C) unstained and (D) stained populations of cells at day 14 of astrocytic differentiation. (E) Quantification of flow cytometric analysis of GFAP expression in differentiating cells (n=3). (F) Representative histogram showing increase in GFAP expression at day 14 compared to day 0 of differentiation. FNSC: Fetal neural stem cells. GFAP: Glial fibrillary acidic protein. Data represented as Mean±SD. * p-value <0.05 **p-value <0.01

### Expression of NSC-marker Nestin and neuronal marker Doublecortin by differentiating cells

Differentiating cells were also evaluated for the expression of the neural stem cell marker, Nestin, at different time points during differentiation, using quantitative PCR. Nestin was found to decrease as differentiation progressed, with a fold change of 0.37 ± 0.25 at day 14 (*p* = 0.18) (Fig. 2E). A significant difference was observed among all the groups (*p* = 0.0432, Kruskal-Wallis test), although significance was not achieved between any two individual groups. Neuronal differentiation was ruled out by evaluating expression of neuronal marker Doublecortin (Fig. 2F) which decreased from day 0 to day 7 (fold change 0.28 ± 0.26, *p* > 0.05) and day 14 (fold change 0.14 ± 0.21, *p*= 0.03).

From these observations, it was evident that human FNSCs successfully differentiated into astrocytes and based on the expression pattern of cell type specific markers, cells at day 14 were subjected to further hypoxia experiments.

### Exposing differentiated astrocytes to hypoxia

Differentiated astrocytes at day 14 with 70-80% confluency were subjected to oxygen concentrations mimicking normoxia (20% and 6%), moderate hypoxia (2%) and severe hypoxia (0.2%) for 48 hours. After exposure to hypoxia, morphological changes did not indicate patterns associated with cell death. Exposure to hypoxia was validated by studying the expression of HIF1α. Western blot analysis showed an increase in the expression of HIF1α protein with increasing grade of hypoxia (Fig. 4A). CA9, VEGF and PGK1 are markers known to be upregulated in hypoxia. Gene expression analysis of CA9 observed in various grades of hypoxia (Fig. 4B) showed a significant increase in moderate (fold change 10.8 ± 10.9, *p* = 0.04) and severe hypoxia (fold change 55.99 ± 53, *p* <0.0001) as compared to normoxia (20% oxygen). A slight increase in CA9 expression was seen in cells exposed to 6% oxygen (fold change 2.1 ± 1.8) but was not found to be statistically significant. VEGF and PGK1 gene expression also increased with hypoxia (data not shown). Samples that showed upregulation of these hypoxia-responsive genes, were analyzed further, along with normoxic controls.

**Figure 4.**
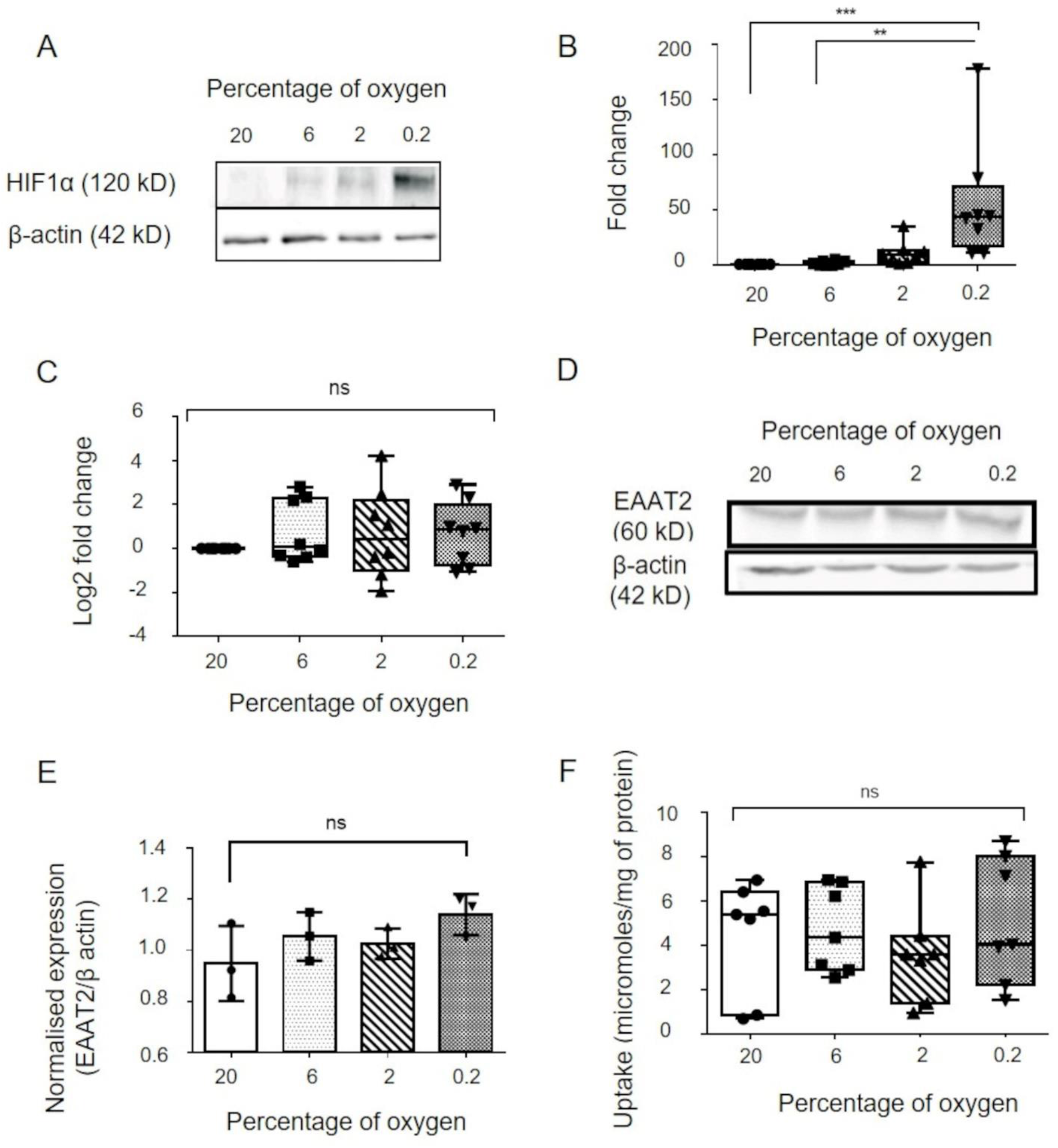
Hypoxia treatment to differentiated astrocytes. (A) Representative western blot image for HIF1α expression in astrocytes exposed to various grades of hypoxia. (B) CA9 and (C) EAAT2 gene expression in astrocytes exposed to various grades of hypoxia as analysed by qPCR (n=8). (D) Representative western blot picture and (E) quantification of western blot data showing expression of EAAT2 in various grades of hypoxia (n=3). (F) Box and whisker plot showing glutamate uptake by astrocytes exposed to different concentrations of oxygen (n=7). CA9: Carbonic anhydrase 9. EAAT2: Excitatory amino acid transporter 2. Data represented as Mean±SD. * p-value <0.05 **p-value < 0.01 ***p-value <0.001 ns: non-significant

### Evaluation of glutamate transporter expression in astrocytes exposed to hypoxia

Expression of the major glial glutamate transporter EAAT2 in astrocytes exposed to hypoxia was studied using qPCR (Fig. 4C) and Western blot (Fig. 4D and 4E). Gene expression analysis of astrocytic EAAT2 at different oxygen concentrations showed log2 fold change (compared to 20% oxygen) values of 0.76 ± 1.4 at 6% oxygen, 0.68 ± 2 at 2 % oxygen and 0.7 ± 1.4 at 0.2 % oxygen. Statistical analysis showed no significant difference between groups. Western blot analysis showed astrocytic expression of EAAT2 (normalised to β-actin) of 0.94 ± 0.14 at 20 % oxygen, 1.05 ± 0.05 at 6% oxygen, 1.02 ± 0.02 at 2% oxygen and 1.14 ± 0.07 at 0.2% oxygen. There was no significant difference detected between groups.

### Evaluation of glutamate uptake in astrocytes exposed to hypoxia

To further delineate the effects of hypoxia on astrocytic glutamate transport, a glutamate uptake assay was performed on astrocytes exposed to hypoxia. Normoxic astrocytes showed a glutamate uptake of 4.4 ± 2.5 micromoles/mg of protein at 20% oxygen and 4.7 ± 1.9 micromoles/mg of protein at 6 % oxygen. Astrocytes maintained glutamate transport at 2% oxygen (glutamate uptake 3.5 ± 2.2 μmoles/mg of protein) and 0.2% oxygen (glutamate uptake 5.1 ± 2.8 μmoles/mg of protein) (Fig. 4F). Statistical analysis showed no significant difference between groups.

## Discussion

Human fetal neural stem cells were isolated from the subventricular region of the brain tissue of aborted fetuses (12-20 weeks gestation) obtained after medical termination of pregnancy. FNSCs showed characteristic features like formation of neuro-spheres in culture, and expression of the neural stem cell marker, Nestin.

Following induction of differentiation in FNSCs, a change in morphology of the cells from small unipolar phenotype of FNSCs, to large, flat polygonal morphology of astrocytes, was observed as differentiation progressed. Although this morphology differs from the classical astrocyte morphology demonstrated *in vivo*, ample evidence exists that *in vitro* cultures that use serum as a differentiation-inducing agent, display fibroblast-like astroglial appearance, similar to that observed in our study (Li et al., 2019). Concomitantly, a gradual rise in GFAP expression peaking at day 14 of differentiation, corresponding with decreasing Nestin expression was observed throughout the course of differentiation. GFAP is the canonical astrocytic marker and has been used for the characterization of astrocytes in numerous studies (Chandrasekharan et al., 2016). Decrease in expression of Nestin observed by us indicates decrease in neural stem cell population as they differentiate into astrocytes. Other studies have observed a similar decline in expression of Nestin following differentiation of neural progenitors into neuronal or glial lineage (Frederiksen and McKay, 1988).

Flow cytometric analysis showed that nearly 95% of cells were expressing GFAP at day 14 of differentiation. Interestingly, 36% of fetal neural stem cells also expressed GFAP. This finding corroborates a previous study, which showed that progenitor cells in adult/embryonic tissue express GFAP (Kim et al., 2018).

In our study, the differentiating cells at day 14 not only expressed GFAP, but also expressed the astrocyte specific glutamate transporter EAAT1 and EAAT2, as well as took up glutamate from the extracellular media, indicating functional activity.

On the basis of the observation that peak GFAP expression was detected at day 14 of differentiation, astrocytes were exposed to different oxygen concentrations at this time-point. Even though cells are usually maintained at 20% oxygen concentration during routine cell culture, various studies reported physiological oxygen concentration in the brain to be much lower, ranging from 1% to 6% (Zhu et al., 2011). Therefore, we exposed astrocytes to oxygen concentrations mimicking normoxia and conditions that may prevail under normoxia *in vivo* (20% and 6%), moderate hypoxia (2%) and severe hypoxia (0.2%) for 48 hours. Cellular response to hypoxia involves the induction of the Hypoxia Inducible Factor 1 alpha (HIF1α) pathway. In our study we saw an increase in the protein expression of HIF1α with an increase in the grade of hypoxia. HIF1α also stimulates the expression of pro-survival and pro-angiogenic molecules such as vascular endothelial growth factor (VEGF), phosphoglycerate kinase (PGK-1) and carbonic anhydrase (CA9), all of which have been demonstrated to be raised following hypoxia (Dengler et al., 2014) and are relatively stable. In our study, we observed that CA9 was robustly increased following exposure of differentiated astrocytes to hypoxia, resulting in a 40-60-fold increase in its expression, consistent with previous studies of hypoxia exposure in astrocytes (Boyd et al., 2017). This was further validated by observing a gradient increase in VEGF and PGK-1 expression with increasing hypoxia (data not shown).

Glutamate is an important excitatory neurotransmitter in the brain, but excessive stimulation of glutamate receptors can lead to neuronal dysfunction and death (Berliocchi et al., 2005). Astrocytes play a key role in preventing glutamate excitotoxicity by taking up excess glutamate from the synapse via the excitatory amino acid transporters (EAATs) out of which EAAT2 is astrocyte-specific and responsible for almost 90% of the glutamate uptake in the forebrain (Kim et al., 2011). Interestingly, we observed that hypoxia exposure did not result in any significant change in the expression of the glutamate transporter EAAT2. Differentiated astrocytes in normoxia showed uptake of glutamate from the extracellular space, which did not change on exposure to hypoxia. This corroborates previous reports that have shown that astrocytes are able to maintain viability in hypoxia, provided energy substrates are present (Almeida et al., 2002). Astrocytic EAAT2 was found to be upregulated in rat models of chronic brain ischemia as well as human tissue (Yatomi et al., 2013) and astrocytes have been shown to continue serving neuroprotective roles in models of intense oxidative stress (Bhatia et al., 2019). Most studies on astroglial glutamate uptake have used rodent models and studies like ours, on primary cultures of non-immortalized human astrocytes, are scarce. To the best of our knowledge, ours is the first study that demonstrates the effects of hypoxia exposure on glutamate uptake in human astrocytes differentiated from fetal neural stem cells.

Even though our findings corroborate studies asserting that astrocytes are resistant to hypoxic conditions, they are in contrast to studies where hypoxic-ischemic injuries to the brain lead to a reduction in glutamate transporter expression (Lukaszevicz et al.,2002). This discrepancy may be due to the added insult of substrate deprivation seen in these studies. Indeed, hypoxia and ischemia have been shown to have differing effects on astrocyte viability and function (Alves et al., 2000). Our findings also differ from those of Dallas et al., (2007) and this disagreement might be due to the different models used, as there are reported differences in the hypoxic sensitivity of mature and developing glia (Hertz et al., 1995).

Some limitations of our study also need to be considered. The time duration of 48 hrs of hypoxia exposure that we have used for our experiments is only a representative selection based on previous work done in our lab. Although astrocytes are also known to release glutamate in response to stimuli such as intracellular calcium changes (Santello and Volterra, 2009), our study does not focus on this aspect of calcium flux on astroglial function.

Our study is novel as we have developed a homogenous population of primary human astrocytes from human FNSCs isolated from aborted fetal brain tissue. The astrocytes thus derived, were characterized with astrocyte-specific markers. Differentiated astrocytes were functional, as indicated by the expression of glutamate transporters and by the uptake of extracellular glutamate by these cells. Moreover, it was observed that these differentiated astrocytes maintained glutamate transporter expression and function, even after exposure to moderate and severe hypoxia. Our unique *in vitro* model of human FNSC derived astrocytes exposed to hypoxic injury, not only corroborates the existing evidence supporting the relative resistance of astrocytes to hypoxic injury, but further substantiates it by demonstrating that astrocytes exposed to hypoxia maintain glutamate uptake. These results thus support the neuroprotective role of astrocytes in hypoxic brain injury, including conditions of severe hypoxia.

## Acknowledgements

The authors are grateful to DBT, Govt. of India, for financial support – extramural research grant (BT/PR21413/MED/122/40/2016). The authors also wish to acknowledge the support of the facilities provided under the Biotechnology Information System Network (BTISNET) grant, DBT, Govt. of India, and Distributed Information Centre at NBRC, Manesar, India.

## Author contributions

### Conception and design of work

Vadanya Shrivastava, Sudip Sen, Pankaj Seth

### Data collection

Vadanya Shrivastava, Devanjan Dey, Chitra Singal, Paritosh Jaiswal, Ankit Singh, JB Sharma

### Data analysis and interpretation

Vadanya Shrivastava, Devanjan Dey, Sudip Sen

### Drafting article

Vadanya Shrivastava, Sudip Sen

### Critical revision of article

Sudip Sen, Subrata Sinha, Pankaj Seth, Parthaprasad Chattopadhyay, JK Palanichamy, JB Sharma.

Final approval of the version to be published: All

## Data accessibility statement

Authors are willing to provide the relevant experimental data upon request.

## Abbreviations used

CA9: carbonic anhydrase 9
EAAT: Excitatory amino acid transporter
FNSC: Fetal neural stem cell
GAPDH: Glyceraldehyde 3 phosphate dehydrogenase
GFAP: Glial fibrillary acidic protein
HIF1α: Hypoxia inducible factor 1 alpha
MEM: Minimum essential media
PGK: Phoshoglycerate kinase
VEGF: Vascular endothelial growth factor

